# Transcriptomic profile of embryoid bodies under hypoxia at single cell level

**DOI:** 10.1101/2025.10.29.685315

**Authors:** Bárbara Acosta-Iborra, Yosra Berrouayel, Laura Puente-Santamaría, Luis del Peso, Benilde Jiménez

## Abstract

Oxygen availability is a key regulator of cellular physiology and hypoxia plays a central role driving vasculogenesis and angiogenesis during development. While bulk transcriptomics has revealed important oxygen-regulated gene networks, such approaches cannot resolve the cellular heterogeneity and lineage dynamics characteristic of early differentiation. To address this, we generated a single-cell transcriptomic dataset from murine embryoid bodies, a widely used in vitro model of early embryonic development, cultured 8 or 10 days under hypoxic (1% O_2_) or normoxic (21% O_2_) conditions for the final 16 or 48 hours of differentiation. This resource enables detailed exploration of how oxygen availability influences lineage specification, vascular and hematopoietic development, and cellular heterogeneity during early differentiation. Beyond developmental biology, the dataset provides a valuable reference for comparative studies of hypoxia responses, benchmarking of single-cell analysis methods, and integrative investigations into oxygen signaling across diverse biological systems.

## Data Description

### Context

Oxygen is a central regulator of cellular metabolism, with hypoxia playing a crucial role in tissue formation and vascular growth during embryogenesis [1]. Here, we present single-cell transcriptomic profiles of mouse embryoid bodies differentiated under controlled oxygen conditions. The dataset comprises thousands of cells collected at two differentiation stages, following either short or prolonged hypoxic exposure, enabling direct comparisons across four experimental conditions. These profiles capture a broad spectrum of developmental states, from progenitors to more differentiated lineages, providing a detailed view of how oxygen availability shapes lineage specification, vascular and hematopoietic development, and cellular heterogeneity. The original aim included examining the generation and expansion of endothelial cells under hypoxia. Although mature endothelial cells were relatively rare in these samples, the dataset captures a broad spectrum of other cell populations, including mesodermal progenitors, endodermal precursors, and more differentiated lineages. Beyond its immediate biological insights, the dataset offers a valuable resource for comparative analyses of differentiation protocols and benchmarking of single-cell computational tools. By making these data openly available, we provide a dataset relevant to research into early cell fate decisions and in vitro models of development.

## Methods

### a) Sampling strategy

Murine embryoid body (EB) samples were generated from the 129 SvJ R1 wild-type mouse embryonic stem cell line (R1 mESCs; ATCC SCRC-1011, RRID:CVCL_2169). Cells were cultured and differentiated as described below. For hypoxic and normoxic comparisons, EBs were maintained in either 1% O_2_ or 21% O_2_ for the last 16 or 48 hours of differentiation, resulting in four experimental conditions in total (Figure 1).

**Figure 1.**
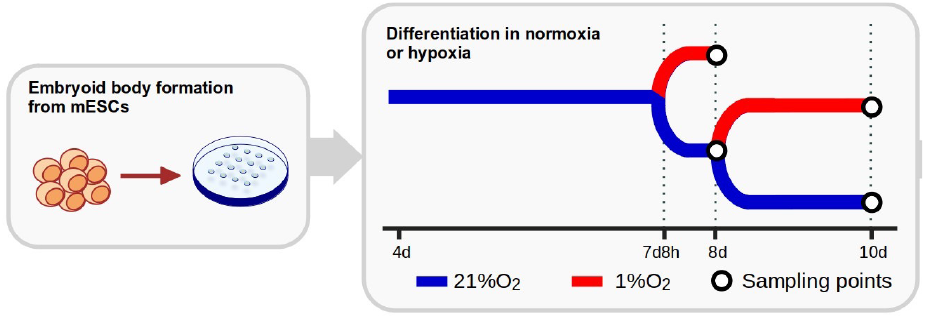
Experimental design. Schematic representation of the experimental design. Mouse embryonic stem cells were differentiated into embryoid bodies for 8 or 10 days under normoxic or hypoxic conditions, with hypoxia applied for 16 h (day 8) or 48 h (day 10).

### b) Steps

- Cell culture and differentiation protocol. R1 mESCs were cultured on a feeder layer of mitomycin C–inactivated mouse embryonic fibroblasts (MEFs; Sigma, M4287) in mESC medium composed of Dulbecco’s modified Eagle’s medium (DMEM; Gibco, 41966052) supplemented with 50 U/mL penicillin and 50 µg/mL streptomycin (Gibco, 15140122), 2 mM GlutaMAX (Sigma, G8541), non-essential amino acids (Thermo Fisher Scientific, 11140050), 0.1 mM 2-mercaptoethanol (Thermo Fisher Scientific, 21985023), 1000 U/mL ESGRO leukemia inhibitory factor (LIF; Chemicon, ESG1106), and 15% (v/v) fetal bovine serum (FBS Biowest, S1300; pre-tested for mESC culture). Medium was refreshed daily, and cells were passaged at a 1:10 ratio every other day. For EB generation, MEFs were separated from mESCs by differential adhesion to 1% gelatin in PBS for 30 min at 37 °C. mESCs were then aggregated in hanging drops (1200 cells per 20 µL drop) and cultured in mESC medium lacking LIF for 4 days. On day 4, R1-derived EBs were plated on 1% gelatin-coated dishes (28 EBs per p60 dish) to induce two-dimensional differentiation. Differentiation proceeded for a total of 8 or 10 days, with subsets of EBs transferred to hypoxia (1% O_2_) or maintained under normoxia (21% O_2_) for the final 16 or 48 h.
- Single-cell dissociation and library preparation. EBs were dissociated to single cells by incubation with Accumax (1 mL per p60 dish; Innovative Cell Technologies) for 30 min at 37 °C. Cells were collected by centrifugation (400 g, 5 min, 4 °C), filtered through a 70 µm strainer (Falcon, 352350), and further dissociated by gentle pipetting with wide-bore tips. Single-cell suspensions were processed for single-cell RNA sequencing (scRNA-seq) using the Chromium Single Cell 3’ Library & Gel Bead Kit (10x Genomics) following the manufacturer’s protocol (10x Genomics, Pleasanton, CA, USA). Libraries were sequenced on an Illumina platform according to 10x Genomics recommendations.
- Data processing and analysis. Raw sequencing reads were processed using Cell Ranger v7.1.0 (10x Genomics; https://support.10xgenomics.com/single-cell-gene-expression/software/pipelines/latest/what-is-cell-ranger). Reads were demultiplexed, aligned to the mouse reference genome (mm10), and gene–cell count matrices were generated.

## Data Validation and Quality Control

Previous studies have shown that hypoxia simultaneously promotes angiogenesis and induces cell cycle arrest in endothelial cells, reflecting an adaptive balance between vascular expansion and reduced proliferation of mature endothelial populations [2]. To confirm that our system recapitulated these established biological responses, we performed several validation experiments prior to sequencing (Figure 2). Flow cytometry analysis of EBs at day 9 revealed an increased proportion of CD31^+^ and CD144^+^ endothelial populations under hypoxic conditions compared to normoxia, indicating enhanced endothelial differentiation. Consistently, immunofluorescence staining of CD31 at day 10 demonstrated more extensive vascular-like networks in hypoxia, with quantitative measurements showing significant increases in branch number, branch length, and total network complexity. In parallel, EdU incorporation assays revealed that hypoxia markedly reduced the proportion of cells in S-phase (Figure 3), confirming cell cycle arrest. Together, these results demonstrate that the EBs responded to hypoxia in a manner consistent with published findings and validate the biological robustness of the dataset.

**Figure 2.**
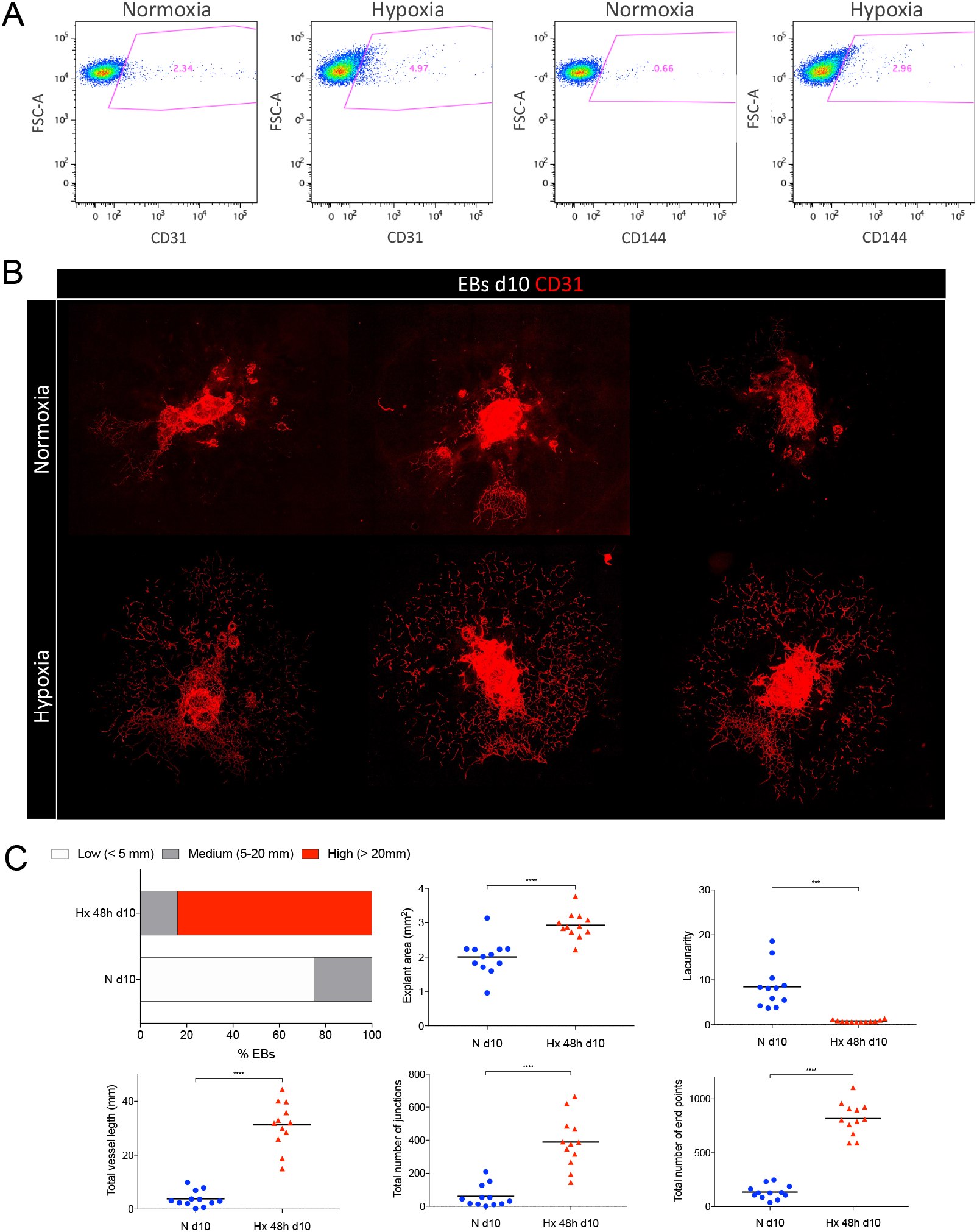
Validation of hypoxia-induced angiogenic responses in embryoid bodies. (A) Flow cytometry plots of day 9 EBs showing CD31^+^ and CD144^+^ endothelial populations under normoxia or hypoxia. (B) Representative immunofluorescence images of CD31^+^ vascular networks in day 10 EBs cultured under normoxia or hypoxia. (C) Quantification of angiogenic parameters with AngioTool analysis software. The upper-left plot shows the percentage of EBs with low (<2.5 mm length, white), medium (2.5-20 mm length, grey) or high (>20 mm length, red) angiogenesis is represented for each experimental condition. A Chi-square analysis showed that the distribution of vessel length was significantly different among conditions (X^2^ =161.2, P<0.0001). The remaining plots show individual AngioTool parameters for each EB: Explant area (mm^2^), Lacunarity, Total vessel length (mm), Total number of junctions and total number of end points. Statistical significance was determined by one-way ANOVA using Kruskal-Wallis post-test (**P <.01, ***P <.001, ****P <.0001)

**Figure 3.**
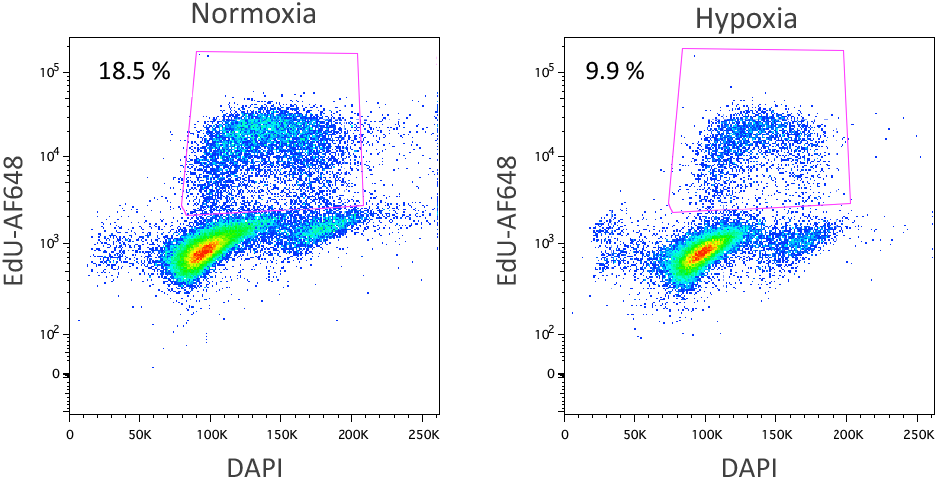
Hypoxia decreases the percentage of endothelial cells in the S-phase of the cell cycle. Representative plot of EdU-Alexa 647 vs DNA content by propidium iodide staining (PI) of day 10 EBs (normoxia or last 24h in hypoxia). Percentage of cells in G0/G1, S, and G2/M phases of the cell cycle is shown inside the corresponding gating regions (magenta lines).

To assess the technical quality of the single-cell RNA-seq dataset, we examined sequencing and mapping metrics across the four experimental conditions. A summary of key quality control statistics is provided in Table 1. Across all samples, high proportions of valid barcodes (>97%) and base quality scores (Q30 >90% across barcode, RNA, and UMI reads) were observed, indicating robust library preparation and sequencing performance. The fraction of reads confidently mapped to the mouse genome (mm10) was consistently >92%, with 70–74% of reads aligning to exonic regions, consistent with high-quality transcript capture. Sequencing saturation ranged from 26.8% to 32.7%, reflecting adequate depth for transcriptome coverage across conditions. The number of estimated cells recovered per sample ranged from 2802 to 4746, with median genes detected per cell between 3104 and 5294, and median unique molecular identifiers (UMIs) per cell ranging from 12,512 to 22,208. These metrics are in line with expected values for 10x Genomics scRNA-seq datasets and confirm successful cell capture and gene expression profiling.

**Table 1.** Summary of sequencing statistics across conditions (Normoxia vs Hypoxia at 8 and 10 days).

| Metric | Normoxia 8d | Hypoxia 8d | Normoxia 10d | Hypoxia 10d |
| --- | --- | --- | --- | --- |
| Estimated Number of Cells | 3374 | 3489 | 2802 | 4746 |
| Mean Reads per Cell | 50 502 | 43 010 | 48 338 | 31 144 |
| Median Genes per Cell | 5294 | 4587 | 4800 | 3104 |
| Number of Reads | 170 394 384 | 150 063 590 | 135 442 885 | 147 809 980 |
| Fraction Reads in Cells (%) | 96.0 | 94.7 | 92.3 | 92.1 |
| Total Genes Detected | 25 219 | 24 024 | 23 968 | 23 695 |
| Median UMI Counts per Cell | 22 208 | 18 349 | 19 825 | 12 512 |
| Valid Barcodes (%) | 97.5 | 97.7 | 97.4 | 97.9 |
| Sequencing Saturation (%) | 30.3 | 28.5 | 32.7 | 26.8 |
| Q30 Bases in Barcode (%) | 94.7 | 94.5 | 94.7 | 94.7 |
| Q30 Bases in RNA Read (%) | 90.5 | 90.4 | 91.4 | 91.4 |
| Q30 Bases in UMI (%) | 94.6 | 94.6 | 94.8 | 94.7 |
| Reads Mapped to Genome (%) | 94.9 | 95.9 | 95.7 | 96.1 |
| Reads Mapped Confidently to Genome (%) | 90.9 | 92.4 | 92.2 | 92.3 |
| Reads Mapped Confidently to Intergenic Regions (%) | 3.1 | 2.4 | 3.2 | 2.7 |
| Reads Mapped Confidently to Intronic Regions (%) | 16.4 | 16.7 | 18.1 | 16.2 |
| Reads Mapped Confidently to Exonic Regions (%) | 71.3 | 73.3 | 70.9 | 73.4 |
| Reads Mapped Confidently to Transcriptome (%) | 81.3 | 83.9 | 81.5 | 83.7 |
| Reads Mapped Antisense to Gene (%) | 6.1 | 5.7 | 7.1 | 5.6 |

Together, these QC results demonstrate that the dataset is technically robust and suitable for downstream analyses, including dimensionality reduction, clustering, and differential expression. In addition, the dataset has been analyzed in a preprint describing hypoxia-induced effects during embryoid body differentiation [3], further supporting the validity and utility of the resource.

## Reuse Potential

This dataset can serve as a valuable reference for multiple applications. It enables comparative studies of early differentiation under varying oxygen conditions and benchmarking of single-cell analysis pipelines. The resource is suitable for integration with other datasets on EB differentiation, enhancing meta-analyses of developmental processes. Furthermore, it may be used in training or testing computational models for cell type classification and transcriptional heterogeneity. Beyond developmental biology, the data may also support broader research on hypoxia-related mechanisms relevant to disease, regeneration, and cellular stress responses.

## List of abbreviations

EB: Embryoid body
GEO: Gene Expression Omnibus
scRNA-seq: Single-cell RNA sequencing
UMI: Unique molecular identifier

## Availability of supporting data and materials

The dataset supporting the results of this article is available in the Gene Expression Omnibus (GEO) repository under accession number GSE226454.

## Declarations

### Ethical Approval

Not applicable.

### Consent for publication

Not applicable.

### Author’s Contributions

Study conceptualization: L.P., B.J.; experimental design: B.J., L.P., B.A.I.; data generation: B.A.I., B.J.; bionformatic analysis: L.P.S, Y.B.; writing original draft: Y.B.; funding acquisition: L.P., B.J.

## Acknowledgements

Please acknowledge anyone who contributed towards the article who does not meet the criteria for authorship including anyone who provided professional writing services or materials.

Authors should obtain permission to acknowledge from all those mentioned in the Acknowledgements section. If you do not have anyone to acknowledge, please write “Not applicable” in this section.

See our [editorial](https://doi.org/10.1093/gigascience/giy122) for a more explanation of acknowledgements and authorship criteria.

